# Fungal-bacterial interaction selects for quorum sensing mutants and a metabolic shift towards the production of natural antifungal compounds

**DOI:** 10.1101/376590

**Authors:** Andrea G. Albarracín Orio, Daniel Petras, Romina A. Tobares, Alexander A. Aksenov, Mingxun Wang, Florencia Juncosa, Pamela Sayago, Alejandro J. Moyano, Pieter C. Dorrestein, Andrea M. Smania, Daniel A. Ducasse

## Abstract

Environmental species of bacteria and fungi coexist and interact showing antagonistic and mutualistic behaviors, mediated by exchange of small diffusible metabolites, driving microbial adaptation to complex communal lifestyles^1^. Here we show that a wild *Bacilus subtilis* strain undergoes heritable phenotypic variation following interaction with the soil fungal pathogen *Setophoma terrestris* (ST) in co-culture. Metabolomics analysis revealed a differential profile in *B. subtilis* before (pre-ST) and after (post-ST) interacting with the fungus, which paradoxically involved the absence of lipopeptides surfactin and plipastatin and yet acquired antifungal activity in post-ST variants. Metabolic changes were also observed in the profile of volatile compounds, with 2-heptanone and 2-octanone being the most discriminating metabolites present at higher concentrations in post-ST during its interaction with the fungus. Most strikingly, both ketones showed strong antifungal activity against *S. terrestris*, which was lost with the addition of exogenous surfactin to the medium. Whole-genome analyses showed that mutations in the *comA* and *comP* genes of the ComQPXA quorum-sensing system, constituted the genetic bases of post-ST conversion, which allowed the concomitant production of ketones and elimination of surfactin. These findings suggest that mutations in ComQXPA stably rewired *B. subtilis* metabolism towards the depletion of surfactins and the production of antifungal compounds during its antagonistic interaction with *S. terrestris*.

---

The great diversity of soil microbes leads to extensive interspecies interactions^1^. Antagonistic and mutualistic behaviors, mediated by exchange of small diffusible secondary metabolites, facilitate microbial adaptation to complex communal lifestyles. *B. subtilis* secretes numerous metabolites, of which many play dual roles as antimicrobial compounds and signaling molecules, participating in processes such as regulation of development, biofilm formation, and inhibition of virulence factors released by competitors^2, 3, 4^. Among them, the cyclic lipopetides surfactin and plipastatin are widely recognized as the main antimicrobials that can be secreted in biologically relevant amounts^5, 6^. Surfactin displays hemolytic, antiviral, antimycoplasma and antibacterial activities, and is essential for biofilm formation, motility and general medium/niche colonization capacity in *B. subtilis*^7, 8, 9, 10^. Thus, surfactin production is regulated by quorum sensing and is activated at the end of exponential growth phase^11^. On the other hand, plipastatin, a member of the fengycin family, displays strong, specific activity against filamentous fung^2, 12, 13, 14^.

We previously isolated a strain of *B. subtilis* (referred to as Bs ALBA01) from the rhizosphere of onion plants that inhibited *in vitro* growth of the soil fungus *Setophoma terrestris*, a major plant pathogen that affects many economically important vegetable crops^15, 16^. After 15 days of interaction with the soil fungal pathogen *S. terrestris* in co-culture assays, *B. subtilis* acquired strong antifungal activity. Such activity was mediated by secreted factors, since bacterial cell-free supernatants displayed fungal growth inhibition^15^ (Supplementary Fig. 1a). These Bs ALBA01 variants were termed post-ST, to distinguish them from pre-ST variants grown without any contact with the fungus and whose cell-free supernatants displayed no antifungal activity^15^ (Supplementary Fig. 1a). Both variants showed clear phenotypic differences, with post-ST variants showing greater degrees of roughness and wrinkled colony phenotype relative to pre-ST variants as well as thicker and more structured pellicles associated with robust biofilm formation (Supplementary Fig. 1a).

Importantly, all post-ST phenotypic traits were maintained when variants were re-inoculated onto fresh growth medium (passaging), indicating that *B. subtilis* experienced genetically stable phenotypic variation upon co-culture with *S. terrestris*.

We next characterized the metabolites profile changes suffered by *B. subtilis* due to conversion to post-ST, which might be responsible for the antagonistic effects observed on *S. terrestris*. Thus, by using non-targeted metabolomic profiling based on ^1^H-NMR spectroscopy, we analyzed cell-free supernatants from pre- and post-ST. Pre-ST samples clustered together and were clearly separated from post-ST samples (Supplementary Fig. 2a and 3). S-line plots in OPLS-DA models showed clearly distinct signals representing discriminant metabolites between the pre- and post-ST sample groups (Supplementary Fig. 2b).

Furthermore, we also compared chemotypes of whole cells in search of significant metabolomic differences between the two variants. Non-targeted high performance liquid chromatography tandem mass spectrometry (HPLC-MS/MS)-based metabolomic analysis revealed separation of global metabolome between the two variants in Principal Coordinates Analysis (PCoA) (Fig. 1a). Surprisingly, the two well-known *Bacillus* lipopeptides, surfactin and plipastatin, were severely reduced in the post-ST variant (Supplementary Fig. 4a and 4b). In fact, random forest analysis showed that they were the most important variables determining differences between pre- and post-ST (Supplementary Fig. 4c). Post-ST showed notable reduction in intensity of surfactin ions and of peaks with masses indicative of plipastatin, identified through spectrum library matching and feature-based molecular networking^17, 18, 19^. Importantly, post-ST clustered together with a *B. subtilis* NCBI 3610 surfactin-defective mutant, confirming the role of surfactin in separation between the groups (Fig. 1a). Molecular networks obtained by analysis of chemical profiles of the two variants indicate reduction in post-ST of not only surfactin and plipastatin derivatives but also related unknown derivatives (Supplementary Fig. 5). We further complemented this analysis with HPLC-MS/MS-based metabolomic analysis of cell-free supernatants of the two variants. Once again, PCoA plots evidenced a discrimination according to the origin of cell-free supernatants, displaying separate clustering of pre-ST vs. post-ST samples (Fig. 1b). Consistently with results of whole-cell analysis, supernatants of post-ST did not show surfactin (Fig. 1c) or plipastatin (Fig. 1d) ions levels comparable to those of pre-ST. Accordingly, post-ST supernatants showed no hemolytic activity (Supplementary Fig. 4d).

**Figure 1.**
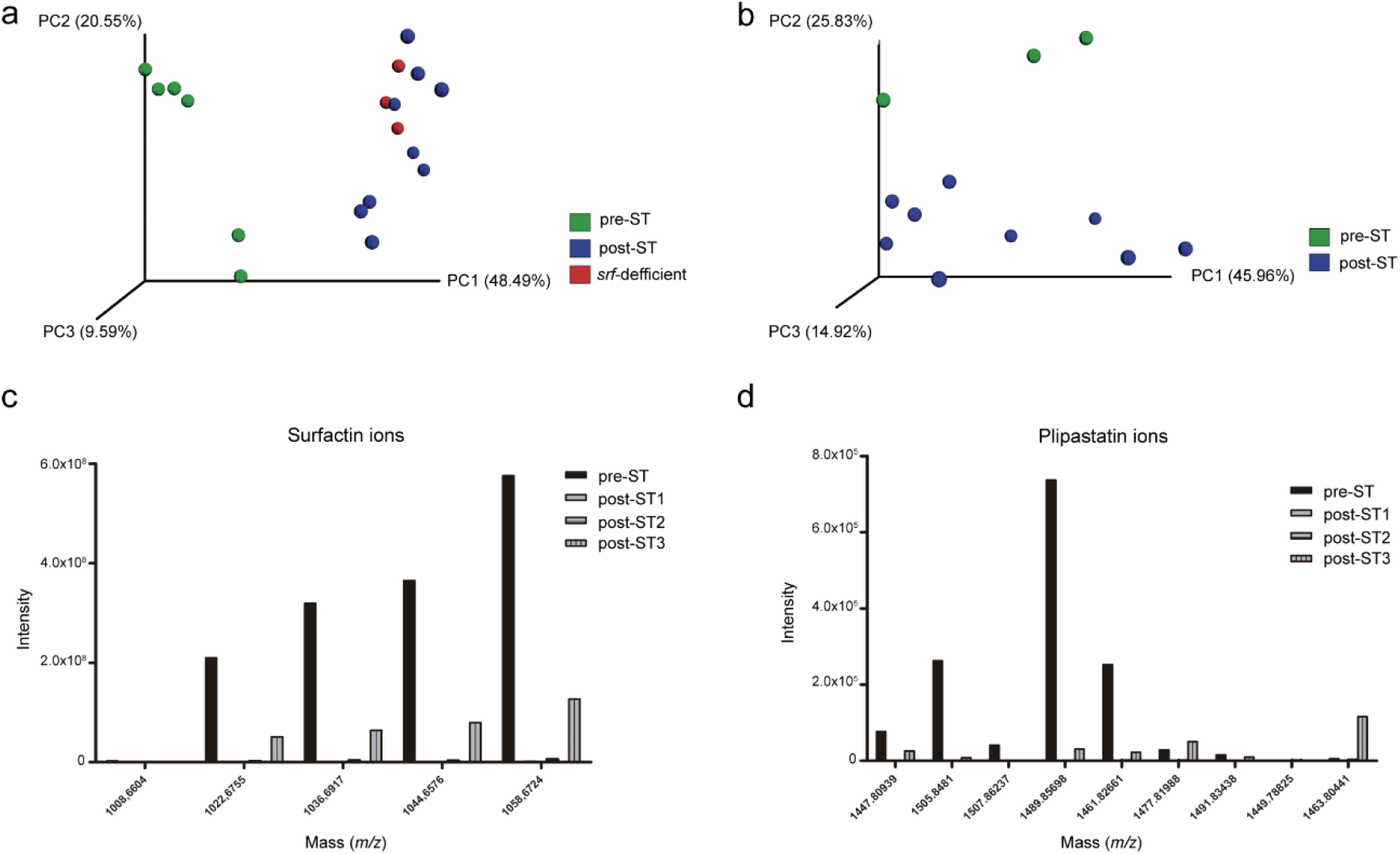
LC-MS/MS-based metabolomic analysis showing chemical signatures that distinguish pre- from post-ST variants. **(a)** PCoA plots showing strong separation between pre- and post-ST. **(b)** PCoA plots showed clear distinctions according to the origin of cell-free supernatants, with separate clustering of pre- vs. post-ST samples. **(c, d)** Intensities of surfactin and plipastatin ions were strongly reduced in in cell-free supernatants of post-ST.

Since the absence of surfactin in post-ST was unexpected, we decided to evaluate the role of surfactin deficiency on *B. subtilis* in this interaction process. To examine the possibility that key genes involved in surfactin production are related to acquisition of antifungal activity by post-ST variants, we generated a surfactin-defective Bs ALBA01 mutant (Δ*srfAA*) by deleting the surfactin synthase encoding gene *srfAA*, essential component of the machinery for non-ribosomal synthesis of surfactin. Anti-*S. terrestris* activity of this mutant was screened upon 15-day of co-culture by obtaining and testing samples before and after co-culture. Interestingly, the absence of surfactin resulted in the acquisition of equivalent mycelial growth-inhibitory activity to that observed in post-ST. In fact, pre- and post-ST supernatants of surfactin-defective Δ*srfAA* showed no notable differences in antifungal activity (Supplementary Fig. 6). Thus, inability to produce/release surfactin resulted in enhanced anti-*S. terrestris* activity as well as complete suppression of ST-driven transformation to post-ST phenotype. On the other hand, unlike the post-ST variants and in agreement with previous reports^20^, the Δ*srfAA* strain lost the capacity to form biofilms.

Given that metabolomics profiles from HPLC-MS/MS showed clear differences, it was intriguing that the main difference in the metabolome of post-ST variants was the suppression of lipopeptides surfactin and plipistatin, whereas no candidate antifungal metabolites showed significantly higher abundances. However, our HPLC-MS/MS analysis was biased towards a limited area of the chemical space, and was unable to detect small primary metabolites and volatile compounds. In order to expand our search to small molecules, we analyzed the volatile compounds profile of the variants and their interaction with *S. terrestris*. Thus, we adapted the growth of co-cultures to glass vials suitable for headspace gas chromatography-mass spectrometry (GC-MS) analysis using solid phase microextraction (SPME). From partial least squares discriminant analysis (PLS-DA), we found that 2-heptanone was the most discriminating compound produced in greater amount post-mutation only by *B. subtilis*, and not by the fungus. Interestingly, by performing a GC data network analysis^21^ within the Global Natural Products Social (GNPS) platform^18^, we could discriminate a cluster composed of a family of ketones, remarkably, all 2-ketones (Fig. 2a). Several of these ketones were the also among top discriminant features between pre-ST and post-ST variants and *S. terrestris* (Fig 2b). Interestingly, individually cultured pre-ST variants showed high levels of 2-ketones (Fig. 2b). However, co-culture with *S. terrestris* was able to induce a steep drop in 2-ketones levels in pre-ST, but remarkably post-ST variant retained the ability to produce them. Thus, we wondered whether 2-heptanone and a representative compound of the ketone cluster, 2-octanone, were sufficient to exert antifungal activity on *S. terrestris*. To the best of our knowledge, there is no specific genic pathway described in *B. subtilis* responsible for the synthesis of 2-ketones and thus no 2-heptanone-nor 2-octanone-deficient mutants of *B. subtilis* can be achieved. Therefore, we decided to test the antifungal effect of either 2-ketones on *S. terrestris*. The activity of both compounds were tested in bioassays by culturing the fungus on PDA dishes containing a filter paper disc where different concentrations of each purified compound were loaded. The growth of *S. terrestris* was gradually reduced as the concentration of 2-heptanone and 2-octanone increased, and the fungal inhibition was maximum at concentrations of 0.02 M and 0.006 M, respectively (Fig. 2c). These results indicate that 2-heptanone and 2-octanone were involved in the antagonistic process against *S. terrestris*. Considering our HPLC-MS/MS results, the antifungal behavior of the Δ*srfAA* mutant strain described above and the lethal effect of both volatiles 2-heptanone and 2-octanone on *S. terrestris*, we next explored whether the presence of surfactin was involved in the lack of antifungal activity characteristically observed in cell-free supernatants of pre-ST variants. If this were the case, then surfactin depletion would be a functional trait of post-ST variants necessary to achieve antifungal capacity. To evaluate this, we added exogenous purified surfactin to cell-free supernatant of post-ST and tested the anti-*Setophoma* activity of this combination. The exogenous surfactin partially suppressed the antifungal activity of post-ST supernatants (Figure 3a). Furthermore, in order to distinguish if this suppressor effect was the outcome of surfactin directly interfering with the activity of antifungal compounds, we tested if surfactin was able to decrease the anti-*Setophoma* activities of 2-heptanone and 2-octanone. The exogenous addition of surfactin protected *S. terrestris* from the antifungal activity of both, 2-heptanone and 2-octanone (Fig. 3b and Supplementary Fig. 7). Although these results are intriguing, previous reports proposed that surfactins could interfere with the activity of other lipopeptides with documented antifungal activity, e.g., fengycins/plipastatins and iturins^3, 22^.

**Figure 2.**
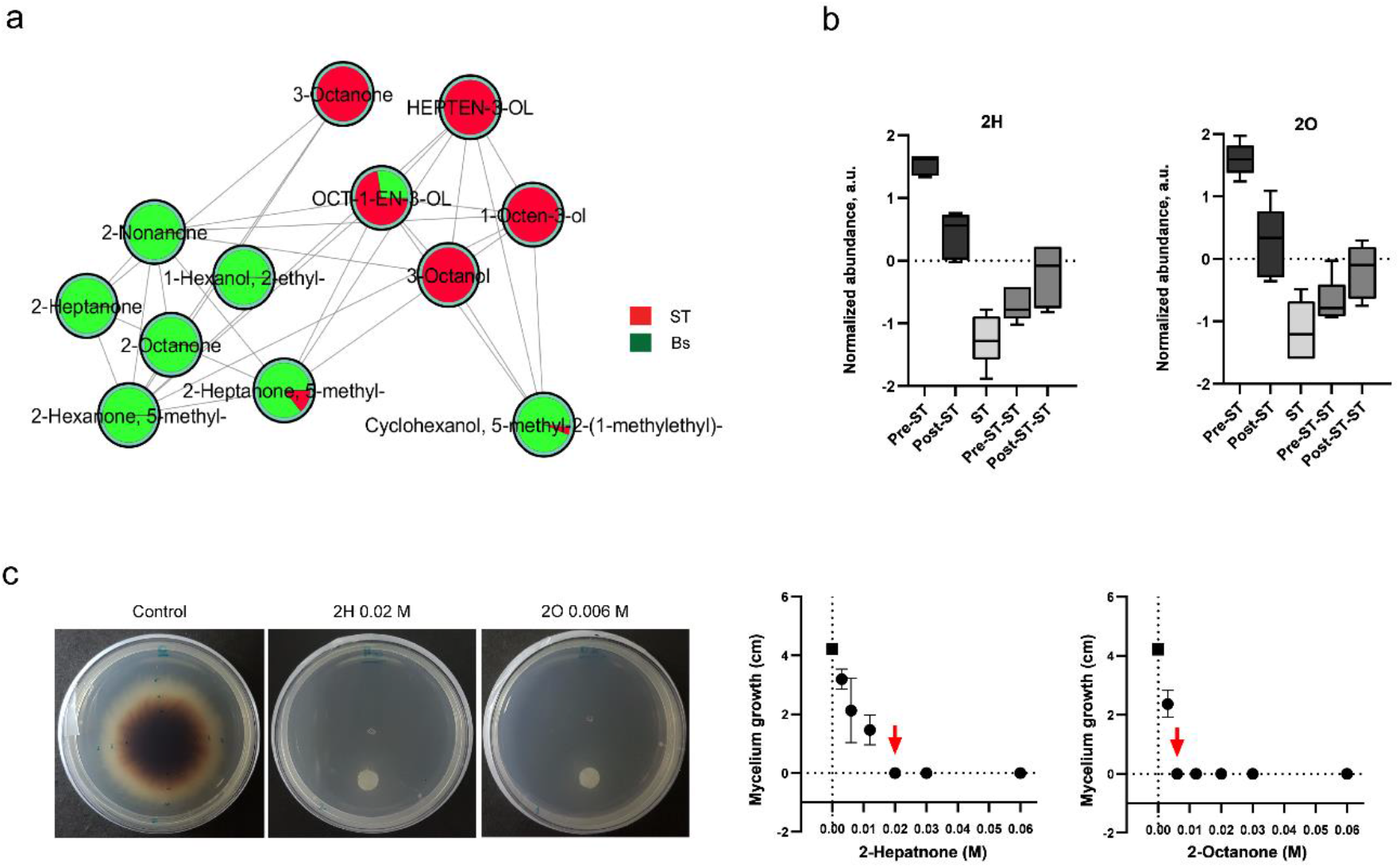
Volatile compounds change due to interaction between *B. subtilis* and *S. terrestris*; 2-ketones induce fungal growth inhibition. **(a)** GC network analysis revealed that Bs ALBA01 produces and releases a family of 2-ketones compounds, which arise as the most important volatile metabolites produced only by the bacterium. **(b)** Interaction with the fungus (ST) generates a steep drop of 2-heptanone (2H) and 2-octanone (2O) levels in pre-ST but not in post-ST. Both, 2H and 2O are the most discriminating features between pre- and post-ST during co-culture with ST. Boxplots show compound abundances, normalized by quantile normalization and auto-scaled (zero-centered and divided by standard deviation). **(c)** Growth inhibition of *S. terrestris* at day 7 after inoculation of 0.02 M 2H and 0.006 M 2O on a filter paper disc placed on PDA dishes. Graphs showing the gradual reduction in mycelium growth with increasing concentrations of 2-ketones. Data shown are mean values of mycelial growth from three independent replicate experiments; red arrow indicates lethal concentration of 2H and 2O at day 7 after inoculation; black square symbol indicates fungal control growth without ketones.

**Figure 3.**
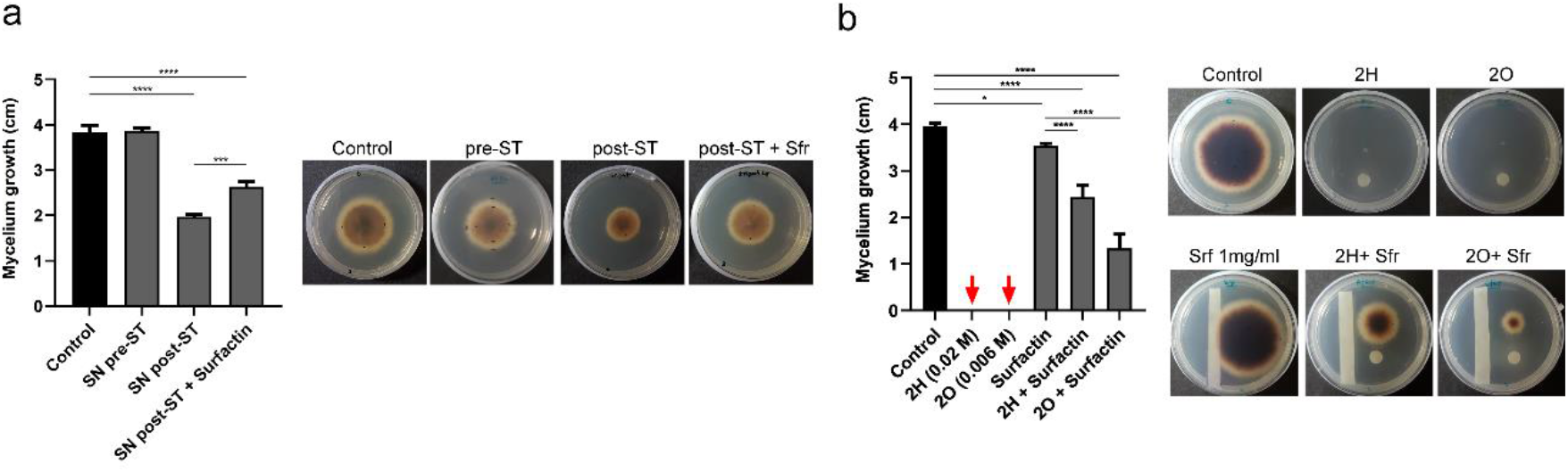
Surfactin interferes with the anti-*S. terrestris* activity of 2-ketones. **(a)** Suppressor effect of exogenous surfactin (1mg/ml) on the antifungal activity of cell-free supernatants of post-ST observed at day 5 after inoculation. **(b)** Suppressor effect of a filter paper strip imbibed with surfactin (1mg/ml) on the antifungal activity of 2H 0.02 M and on 2O 0.006 M, 7 days after inoculation. Data shown are mean values of mycelial growth from three independent replicate experiments; red arrow indicates lethal concentration of ketones at day 7.

Finally, we sequenced whole genomes of three post-ST variants (post-ST1, post-ST2, post-ST3; Supplementary Fig. 8) derived independently from Bs ALBA01, which was considered as pre-ST ancestor and used as reference. Reads from these three variants were aligned against Bs ALBA01 genome^16^ (NCBI Bioproject PRJNA316980) to assess genetic changes accumulated in post-ST during interaction with the fungus. We found one to three different mutations in coding sequence regions for each of the variants (Supplementary Table 1). Interestingly, each variant had at least one loss-of-function mutation in genes of the ComQXPA quorum-sensing system. The post-ST1 and post-ST2 variants had a frameshift mutation in coding sequence of the *comA* gene, while post-ST3 variant harbored a 100 bp deletion in the kinase-encoding gene *comP.* Mutations in *comA* and *comP* were further confirmed by PCR amplification and sequencing.

Then, we investigated the extent to which *com* genes in Bs ALBA01 were convergently mutated upon interaction with *S. terrestris* by performing whole-genome sequencing of a pool of 15 post-ST variants obtained independently following interaction with the fungus in co-culture (Pool-seq; see Methods). To identify allelic variations in the pool, we mapped sequence reads to the *B. subtilis* ALBA01 reference genome, and then performed variant calling (see Supplementary Methods). Of note, six different loss-of-function mutations were identified in the *comQXPA* operon within the pool of post-ST variants, revealing marked convergence of mutation within these genes (Supplementary Table 1). These findings demonstrate the occurrence of mutation-based phenotypic changes in post-ST variants and support the notion that interaction of Bs ALBA01 with *S. terrestris* induces metabolomic changes promoting bacterial antifungal activity, which involves the ComQXPA quorum-sensing system. In addition, they suggest that single mutations in one of the ComQXPA genic components provided pleiotropic effects for the adaptation of *B. subtilis* to antagonistic interactions.

The ComQXPA system has been reported to be required for transcription of numerous genes involved in competence development, antibiotic production, exopolysaccharide production, degradative enzyme production and transport, and fatty acid metabolism^23,24^. In fact, over 10% of the *B. subtilis* genome is controlled by the ComQXPA quorum-sensing system^24^. Notably, ComA activates transcription of the *srfA* operon that governs surfactin production^25, 26^. The ComQXPA system has also been reported to play a role in transcription of regulator DegQ, which controls production of degradative enzymes and is required for plipastatin production^27, 28, 29^. Thus, mutations in ComQXPA are in agreement with the loss of surfactin and plipastatin production observed in post-ST. On the other hand, mutations in ComA could possibly provoke alterations on the general metabolism promoting chemical conditions that favor the biosynthesis of 2-heptanone and 2-octanone, as those produced through the degradation of branched-chain amino acids, leucine, valine and isoleucine, which would be otherwise used for surfactin or lipopetides synthesis. Alternatively, it has also been described that ComA controls the expression of FapR, a transcriptional regulator involved in fatty acid synthesis in *B. subtilis*^24^. In this sense, FapR negatively regulates the expression of at least 10 genes (the *fap* regulon)^30^. Thus, mutations in *comA/comP* could determine an increased fatty acid biosynthesis by downregulating FapR. The release of the *fap* regulon in association to the chemical condition derived from the global change produced by the mutation may be favoring branched-chain fatty acid biosynthesis pathways and/or decarboxilation of intermediates ketoacids that promote the synthesis of 2-ketones. Accordingly, a recent report showed an enhancement of *de novo* fatty acid synthesis in *Bacillus nematocida* B16 during 2-heptanone production^31^.

This work uncovers the ComQXPA quorum sensing system as a novel pathway employed by *B. subtilis* to achieve stable metabolic rewiring which enables the production of extracellular metabolites required to outcompete microbial foes in antagonistic interactions. The results suggest that mutations in ComQXPA provided a metabolic shift that resulted in two major outcomes: *i)* post-ST variants were capable of retaining ability to produce effective levels of antifungal 2-ketones upon interaction with the fungus while *ii)* switching off the production of surfactin, which would otherwise interfere with the antifungal activity of 2-ketones. As a bonus, mutations in ComQXPA also promoted biofilm formation, another important feature to overcome biological interactions in natural habitats such as the root of plants.

Taken together, our findings show that the interaction of *B. subtilis* with *S. terrestris* drives specific mutation-based bacterial transformation that involves quorum-sensing genes and mediates an adaptive process whereby the bacterium restructures its general metabolism and profile of released molecules to respond to and ultimately overcome the fungal pathogen.

## Methods

### Microorganisms and culture conditions

Bacterial and fungal strains used in this study are listed in Supplementary Table 2. Bacterial strains were routinely grown in LB medium (10 g/L tryptone, 5 g/L yeast extract, 10 g/L NaCl) at 30ºC. For biofilm assays, cells were grown in either LB or LB plus glycerol and manganese (LBGM) at 30ºC. When necessary, LB and LBGM media were solidified with 1.5% agar.

### Construction of mutant strains

Integrative plasmid was constructed to disrupt *srfAA* gene. To construct pSG1194-srfAA, an internal fragment *srfAA* from *B. subtilis* ALBA01 were amplified using primer pairs (listed in Supplementary Table 3) containing EcoRI and BamHI restriction sites. PCR products were digested with EcoRI and BamHI and ligated into pSG1194 vector digested with the same enzymes. Ligation mixtures were transformed into heat shock *E. coli* DH5α competent cells. Plasmids were purified, and ALBA01 competent cells were transformed by standard protocols^32^ to generate the mutant strain Δ*srfAA*. For confirmation of gene disruption and selection of single-copy transformants, chloramphenicol-resistant colonies were analyzed by PCR and subjected to Sanger sequencing (Unidad de Genómica, Instituto de Biotecnología, C.I.C.V.yA. Instituto Nacional de Tecnología Agropecuaria (INTA), Buenos Aires, Argentina).

### Whole genome sequencing and analysis

Whole genome sequencing was performed using a paired-end (PE) 2×100 bp library on Illumina Hiseq 1500 system (INDEAR Genome Sequencing facility, Argentina), and *de novo* assembly on A5 pipeline^33^. ALBA01 reads were assembled into 28 scaffolds with average scaffold size 147,128 bp. For annotation of the genome, scaffolds were uploaded to <u>R</u>apid <u>A</u>nnotation using <u>S</u>ubsystem <u>T</u>echnology (RAST) server^34^, and SEED-based method was applied on this server. The resulting assembly was used as reference genome sequence to map reads from post-ST variants using BWA-MEM tool (v. 0.7.5a-r405)^35, 36^. Variants were called using Genome Analysis Toolkit (GATK) HaplotypeCaller ^37^, and single-nucleotide polymorphisms (SNPs) were filtered based on Phred score >99%. SNPs detected from analysis of ancestral reads were excluded as false positives. Integrative Genomics Viewer (IGV)^38, 39^ was used for manual inspection of variants and read alignments. *comA* and *comP* gene mutations were validated to confirm their presence in post-ST variants and absence in pre-ST variants or ALBA01. For this, a 484-bp *comA* fragment was amplified by PCR using oligonucleotide primers FcomA_map and RcomA_map (Supplementary Table 3). To map mutations in *comP*, a PCR fragment was amplified using primer pair FcomP_map/RcomP_map. PCR products were purified with Silica Bead DNA Gel Extraction Kit (Thermo Fisher, USA), subjected to Sanger sequencing, and analyzed using BioEdit^40^.

### Whole-genome sequencing of pools of individuals (Pool-seq) analysis

Fifteen *B. subtilis* post-ST variants were obtained from independent co-culture experiments as described above. Genomic DNA was collected and purified from each clone using Promega Wizard Genomic DNA purification kit as per the manufacturer's protocol. DNA quantity was determined using Qubit Fluorometric Quantitation fluorometer (Thermo Fisher), and appropriate dilutions were mixed together such that each genome was represented equally in the final pool. Libraries were prepared using Nextera XT DNA Library Preparation Kit (Illumina, USA) as per the manufacturer's protocol. Paired-end sequencing was performed on an Illumina MiSeq platform producing 2×150 bp read lengths (INDEAR Genome Sequencing). Analysis of raw data quality was performed with FastQC^41^. Adapter sequences were trimmed using Trimmomatic^42^. Pool-seq reads were mapped against ALBA01 reference genome using BWA ALN^36^. PCR duplicates, multialignment reads, improperly paired aligned reads, and soft-clipped alignments were removed using Samtools^43^. Indels were realigned with GATK^37^. SNP, insertion, and deletion discovery was performed using HaplotypeCaller algorithm^37^ with sample ploidy parameter set to 15. Under this methodology, variants detected by analyzing ALBA01 ancestral reads were excluded as potential false positives. Filter sequence variants was performed by running GATK VariantFiltration with parameters “QD < 2.0 || FS > 60.0 || MQ < 40.0 || MQRankSum < −12.5 || ReadPosRankSum < −8.0” for SNP, and “QD < 2.0 || FS > 200.0 || ReadPosRankSum < −20.0” for indels. Nucleotide sequence accession numbers: sequence of ALBA01 assembled genome was deposited in NCBI database (Bioproject PRJNA316980)^16^. Genome sequence reads from post-ST variants and Pool-seq analysis were deposited in NCBI database (Bioproject PRJNA480851).

### High performance liquid chromatography – tandem Mass Spectrometry (HPLC-MS/MS)

Biological triplicates of *Bacillus subtilis* pre and post-ST variants as well as a *B. subtilis* NCBI 3610 wild-type strain and a *B. subtilis* NCBI 3610 knock-out *srfAA* mutant were cultivated in LB medium at 30 °C at 180 rpm. The cultures were harvested after 18 hours through centrifugation at 7000 rpm, 4 °C for 5 min. Cell metabolism was quenched by resuspending the cell pellet in −80 °C methanol to a final concentration of 500 mg/ml to final volume of 500 μl in a 96 deepwell plate. The samples were then sonicated 10 minutes and the cell debris was pelleted by 10 min centrifugation at 7000 rpm. 450 μl of supernatants were transferred to a new Deepwell plate and dried overnight in a CentriVap Benchtop Vacuum Centrifuge (LabConco). The metabolites were then resuspended in 100 μl 50% methanol, 1% formic acid and submitted for HPLC-MS/MS analysis. Non-targeted HPLC-MS/MS analysis was performed according to Petras *et al.* 2016^44^. Therefore, 5 uL of the samples were injected on a Q-Exactive Quadrupole-Orbitrap mass spectrometer coupled to Vanquish ultra-high performance liquid chromatography (UHPLC) system (Thermo Fisher Scientific, Bremen, Germany). For the LC separation, a C18 core-shell column (Kinetex, 50 × 2 mm, 1.8 um particle size, 100 A pore size, Phenomenex, Torrance, USA) with a flow rate of 0.5 mL/min (Solvent A: H2O + 0.1 % formic acid (FA), Solvent B: Acetonitrile (ACN) + 0.1 % FA) was used. During LC-MS/MS analysis, the compounds were eluted with a linear gradient from 0-0.5 min, 5 % B, 0.5-4 min 5-50 % B, 4-5 min 50-99 % B, flowed by a 2 min washout phase at 99% B and a 2 min re-equilibration phase at 5 % B. For positive mode MS/MS analysis the electrospray ionization (ESI) parameters were set to 35 L/min sheath gas flow, 10 L/min auxiliary gas flow, 2 L/min sweep gas flow and 400 °C auxiliary gas temperature. The spray voltage was set to 3.5 kV and the inlet capillary was set to 250 °C. A S-lens voltage of 50 V was applied. MS/MS product ion spectra were recorded in data dependent acquisition (DDA) mode. Both MS1 survey scans (150-1500 m/z) and up to 5 MS/MS scans per duty cycle were measured with a resolution (R) of 17,500 with 1 micro-scan. The maximum C-trap fill time was set to 100 ms. Quadrupole precursor selection windows were set to 1 m/z. Normalized collision energy was stepwise increased from 20 to 30 to 40 % with z = 1 as default charge state. MS/MS scans were automatically triggered at the apex of chromatographic peaks within 2 to 15 s from their first occurrence. Dynamic exclusion was set to 5s. Ion species with unassigned charge states and isotope peaks were excluded from MS/MS acquisition.

Detailed descriptions of data analysis, MS/MS network analysis and metabolomic studies based on NMR spectroscopy are presented in the Supplementary Information section. MS/MS Data availability: all HPLC-MS/MS data can be found on the Mass spectrometry Interactive Virtual Environment (MassIVE) at https://massive.ucsd.edu/ with the identifier MSV000082081.

### Gas chromatography – Mass Spectrometry (GC-MS) metabolomics

Biological replicates of *Bacillus subtilis* pre and post-ST variants and of the fungus *S. terrestris* were cultivated individually and in co-cultures in 10 ml-borosilicate vials with a screw cap with silicon septum containing three ml of PDA medium for seven days at 30°C and then analyzed with GC-MS to determine the emitted volatile metabolites. After incubation, the capped vials were stored at −80 °C and thawed immediately prior to analysis. The GC-MS analysis was carried out using the Agilent 7200 GC/QTOF equipped with robotic sampler system. The volatiles from the sample were extracted from headspace using Polydimethylsiloxane/Divinylbenzene (PDMS/DVB) df 65 μm Solid Phase Microextraction Fiber (SPME) for 30 minutes at 50 °C. The fiber was then inserted into the injector equipped with Merlin septum heated to 250 °C and the adsorbed compounds were desorbed for 1 minute. The GC protocol analysis included: cryofocusing on the head of the column at −20 °C for 1 min; 115 °C/min oven ramp to 40 °C (hold of 0.1 min), 20 °C/min oven ramp to 300 °C (hold of 0.1 min), ramp to 320 °C and a), and 50 °C/min oven ramp to 320 °C purge the column. The helium carrier gas was set to constant 2 mL/min flow and a splitless injection mode was applied. The scanned *m*/*z* range was 35-400 Th with the acquisition rate of 20 spectra/s. The empty vial blanks were interspersed with the samples to assess background signal. Quality controls of natural mint oil extract were run along with samples before and after the analysis. The GC-MS data were processed with MZmine2 (https://bmcbioinformatics.biomedcentral.com/articles/10.1186/1471-2105-11-395) using ADAP algorithm (https://pubs.acs.org/doi/full/10.1021/acs.jproteome.7b00633?src=recsys) deployed on the ProteoSAFE workflow of the GNPS platform (gnps.ucsd.edu). The parameters were set as indicated in the Supplementary Table 4. Data were uploaded to GNPS and searched against NIST 2017 and WILEY spectral libraries (link to GNPS search results: https://gnps.ucsd.edu/ProteoSAFe/status.jsp?task=9569f1598a414e9a926721942e41ed6c). The GC-MS data are available at the MassIVE depository (massive.ucsd.edu) under ID MSV000083294.

## Acknowledgments

This study was supported by MINCyT-Agencia Nacional de Promoción Científica y Tecnológica (ANPCyT) (grants PICT 2013-2592 to A.G.A.O and PICT 2016-1545 to A.M.S.), U.S. National Science Foundation (NSF) Inspire Track II (grant IOS-1343020), National Institutes of Health (NIH) (grants GMS10RR029121, 5P41GM103484–07), and Deutsche Forschungsgemeinschaft (DFG) (grant PE 2600/1). The authors are grateful to Dr. Zdenek Kamenik, Institute of Microbiology of the Czech Academy of Sciences, for kindly providing ketones compounds, to Consejo Nacional de Investigaciones Científicas y Técnicas (CONICET), Fulbright Commission Argentina and to Dr. S. Anderson for English editing of the manuscript.

## Author contributions

A.G.A.O and A.M.S designed the experiments and supervised the study. A.G.A.O, F.J., and P.S. cultured the strains. A.G.A.O. performed interaction experiments. A.G.A.O, R.A.T., A.J.M. and A.M.S. performed DNA isolation and analyzed bioinformatic data of bacterial genomes. A.G.A.O. analyzed NMR data. A.G.A.O., D.P. and A.A.A. collected MS data. A.G.A.O., D.P., A.A.A., M.W and P.C.D. analyzed metabolomic data and performed molecular networking. A.G.A.O. and A.M.S. integrated all experimental data and made final interpretations. A.G.A.O., A.J.M., A.M.S. and D.A.D. wrote the manuscript. All authors discussed, edited, and approved the finalized manuscript.

