## Supplementary material for "Fungal-bacterial interaction selects for quorum sensing mutants and a metabolic shift towards the production of natural antifungal compounds": SI material

[2]Universidad Nacional de Córdoba. Facultad de Ciencias Químicas. Departamento de Química Biológica Ranwel Caputto. Córdoba, Argentina.

[3]CONICET. Universidad Nacional de Córdoba. Centro de Investigaciones en Química Biológica de Córdoba (CIQUIBIC), Córdoba, Argentina.

[4]Collaborative Mass Spectrometry Innovation Center, University of California, San Diego. La Jolla, CA 92093, USA.

[5]Scripps Institution of Oceanography, University of California San Diego. La Jolla, CA 92093, USA.

[6]Instituto de Patología Vegetal IPAVE-CIAP, Instituto Nacional de Tecnología Agropecuaria. Córdoba, Argentina.

[‡] These authors contributed equally to this work

**Supplementary file includes:**

Supplementary Methods

Supplementary Figures 1 to 8

Supplementary Tables 1 to 4

References

### Supplementary methods

#### Biofilm formation, colony morphology, and swarming motility

For pellicle formation, starting cultures of each strain were grown in 5 ml LB for 8 h at 37°C with agitation. Three µl of the starting culture were used to inoculate 3 ml LBGM in 96-well microtiter plates and in glass tubes, and incubation without agitation was performed at 30°C. To evaluate colony morphology, 2 µl of the starting culture were spotted onto the surface of LB and LBGM 1.2% agar plates, incubated 3-5 days at 30°C, and colonies were analyzed and photographed. To evaluate swarming motility, cells were collected from an overnight colony with a sterile toothpick and inoculated in the center of LB plates with 0.7% agar. Plates were incubated at 37°C and evaluated for colony spread as a function of time.

#### MS/MS data and featured-based molecular networking analysis

Following HPLC-MS/MS data acquisition, raw spectra were converted to .mzXML file format using MSConvert tool (ProteoWizard). MS1 and MS/MS feature extraction was performed with MZmine v. 2.30 [1]. Intensity thresholds 1E5 and 1E3 were used for MS1 and MS/MS spectra, respectively. For MS1 chromatogram building, mass accuracy 10 ppm and minimum peak intensity 5E5 were set. Extracted Ion Chromatograms (XICs) were deconvoluted using baseline cutoff at intensity 1E5. After deconvolution, MS1 features were matched to MS/MS spectra within 0.02 m/z and 0.2 min retention time windows. Isotope peaks were grouped, and features from different samples were aligned with mass tolerance 10 ppm and retention time tolerance 0.1 min. MS1 features without assigned MS2 features were filtered out of the resulting matrix, as were features that did not contain isotope peaks or did not occur in at least three samples. After filtering, gaps in the feature matrix were filled with relaxed retention time tolerance 0.2 min and mass tolerance 10 ppm. The feature table was exported as .csv file and corresponding MS/MS spectra as .mgf. file. Contaminate features observed in Blank samples were filtered, and only those with relative abundance ratio blank to average in the samples <50% were considered for further analysis.

For feature-based molecular networking, the .mgf file was uploaded to GNPS (gnps.ucsd.edu) [2, 3]. For networking, minimum cosine score to define a correlation between spectra was set to 0.7, Precursor Ion Mass Tolerance to 0.01 Da, Fragment Ion Mass Tolerance to 0.01 Da, Minimum Matched Fragment Ions to 4, Minimum Cluster Size to 1 (MS Cluster off), and Library Search Minimum Matched Peaks to 4. When Analog Search was performed, Cosine Score Threshold was 0.7 and Maximum Analog Search Mass Difference was 100. Molecular networks were visualized using software program Cytoscape v. 3.4 [4], and node information was matched with MS1 feature table. Tridimensional PCoA plots of MS1 data were generated by in-house tool ClusterApp using Bray-Curtis distance metric. Resulting scatter plots were visualized on EMPeror. Random forest classification was performed in R.

### $^1$H-NMR spectroscopy-based metabolic profiling of cell-free supernatants of *B. subtilis* pre- and post-ST variants

$^1$H-NMR spectroscopy and multivariate data analysis were performed at PLABEM (Plataforma Argentina de Biología Estructural y Metabolómica; Rosario, Argentina). Samples were prepared for $^1$H-NMR as described in [5]. Briefly, 540 µl of the supernatant were mixed with 60 µl phosphate buffer (pH 7.4) containing sodium 3-trimethylsilyl-(2,2,3,3-$^2H_4$)-1-propionate (TSP) in $D_2O$ (final concentration 0.1 mg/mL). TSP acts as internal chemical shift reference (δ= 0.0), while $D_2O$ provides lock signal for the spectrometer. Samples were stood for 10 min, then centrifuged at 4000 rpm for 10 min to remove any precipitates. 500 µl of centrifuged solution were transferred to NMR tube. A pooled quality control sample (QC (2)) was prepared by mixing equal volumes (100 µL) of all 12 samples.

Spectra were obtained at 300 K using an Avance 600 MHz NMR spectrometer (Bruker BioSpin; Rheinstetten, Germany) equipped with 5-mm TXI probe. One-dimensional $^1$H-

NMR spectra of conditioned culture media were acquired using standard 1-D NOESY pulse sequence (noesygppr1d) with water presaturation [6]. Mixing time was set to 10 ms, data acquisition period to 2.73 s, and relaxation delay to 4 µs. $^1$H-NMR spectra were acquired using 4 dummy scans and 32 scans, with 64K time domain points and spectral window 20 ppm. FIDs were multiplied by an exponential weighting function corresponding to line broadening 0.3 Hz.

Spectroscopic data was processed by MATLAB v. R2015b (MathWorks Inc.; U.S.). Spectra were referenced to TSP at 0.0 ppm, with baseline correction and phasing of spectra performed using in-house software (provided by T. Ebbels and H. Keun, Imperial College, UK). Each spectrum was reduced to a series of integrated regions of equal width (0.04 ppm, standard bucket width). Spectral regions containing no metabolite signals and TSP signal <0.2 ppm, and the interval containing the water signal (between 4.9 and 4.6 ppm) were excluded. Each spectrum was then normalized by probabilistic quotient method [7].

Pre-processed $^1$H-NMR spectral data were imported to SIMCA (v. 14.1, Umetrics AB; Umeå, Sweden) for multivariate data analysis. Principal Component Analysis (PCA) was performed using the Pareto-scaled NMR dataset. Orthogonal partial least squares discriminative analysis (OPLS-DA) was performed to maximize separation between treatment groups. S-line plots (tailored S-plots) [8] useful for NMR data analysis) were generated to visualize differences between classes in OPLS-DA models. Full cross validation (CV) was performed to ensure valid and reliable models and to avoid overfitting [9].

### Supplementary figures and legends

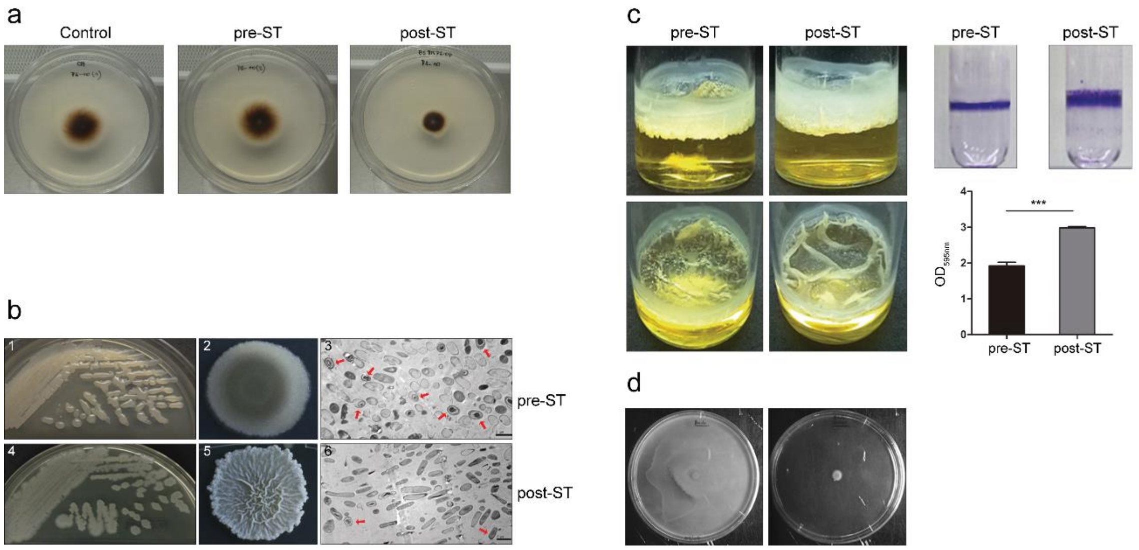


**Supplementary Figure 1. Co-culture of *B. subtilis* with *S. terrestris* induced stable phenotypic changes.**

**(a)** Growth inhibition of *S. terrestris* at day 4 after inoculation. Strong growth inhibition was observed in dishes containing only cell-free supernatants of post-ST. **(b)** Phenotypic changes of Bs ALBA01 resulting from interaction with fungus in LB agar culture. Colonies of post-ST were rougher and more wrinkled than those of pre-ST. Agar plates (panels 1 and 4) and individual colonies (panels 2 and 5) of pre- and post-ST, respectively. Electron microscopy revealed increased numbers of elongated cells and reduced numbers of sporulated cells in post-ST (panel 6) relative to pre-ST (panel 3). Scale bar: 2 µm. Arrows: sporulated cells. **(c)** Post-ST variants in liquid medium developed thick, wrinkled pellicles characteristic of strong biofilm formation. Quantification of biofilm formation on borosilicate surfaces by crystal violet staining revealed greater biofilm formation by post-ST than by pre-ST. Data shown are mean values of measurements of optical density at 595 nm of crystal violet suspensions from three independent replicate experiments. ***, significant difference between values ($p < 0.0001$, Tukey's Multiple Comparison Test). **(d)** Swarming motility assessed on 0.7% agar LB plates. Pre-ST displayed full motility (*i.e.*, covered the entire plate), whereas post-ST showed no swarming motility (*i.e.*, did not extend beyond the inoculation area).

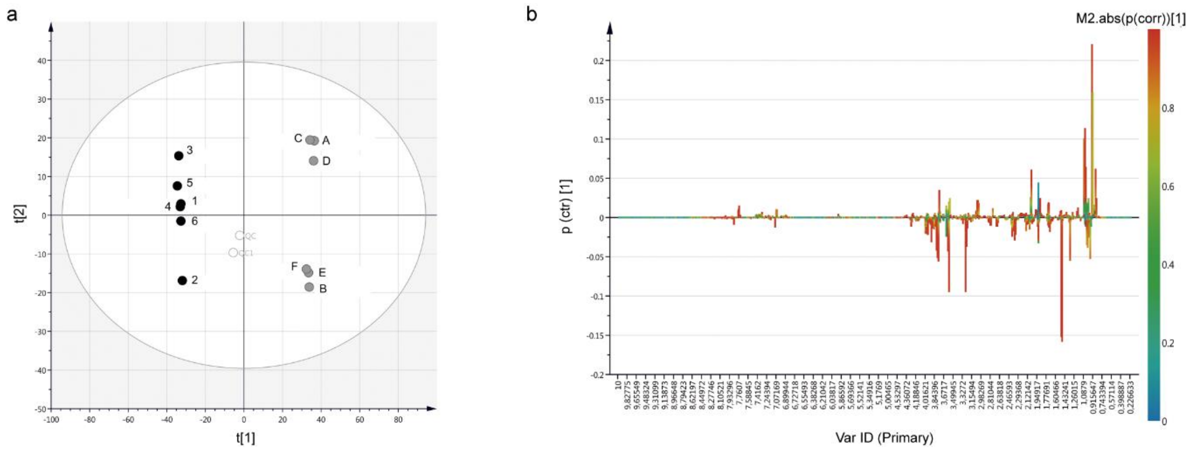


**Supplementary Figure 2. Metabolomic profile variation in *B. subtilis* ALBA01 was observed following *in vitro* interaction with *S. terrestris***

High antifungal activity of cell-free supernatants of post-ST variants appeared to be associated with changes in metabolomic profiles of released compounds, because differences in cell-free supernatants of pre- vs. post-ST variants were detectable using untargeted NMR approach. **(a)** Plot of PCA scores shows clear separation of supernatant samples according to their type (black, pre-ST; gray, post-ST) in the first two principal components derived from $^{1}$H-NMR spectra of *Bacillus*-conditioned media (percent variation in NMR data explained by the model, R2x= 70.4%; percent variation in NMR data predicted by the model from cross-validation, Q2x= 57%, based on 2-components model). White circles: quality control (QC) samples. **(b)** OPLS-DA S-line plots with pairwise comparison of data from NMR spectra from cell-free supernatants of pre- and post-ST. Colors are associated with correlation of metabolites characterized from $^{1}$H-NMR data for the class of interest, using the scale on the right.

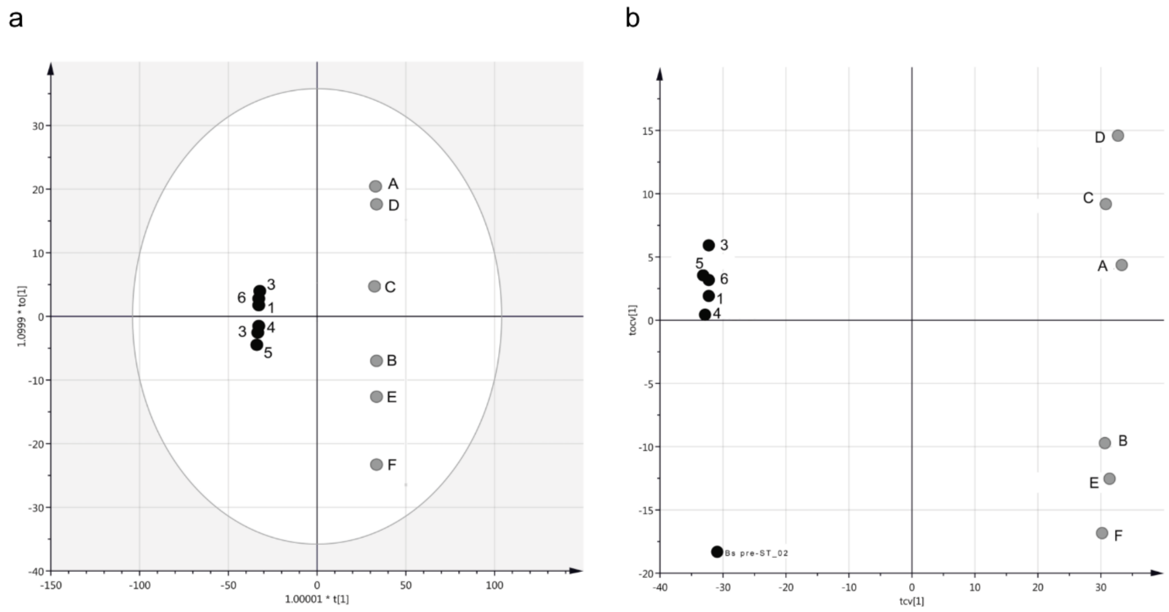


**Supplementary Figure 3. Effect of *S. terrestris* interaction with co-cultured *B. subtilis* on metabolic fingerprints and footprints**

**(a)** Plots of OPLS-DA scores and cross-validated scores for pre-ST (black) and post-ST (gray) samples. This analysis was performed to rule out potential bias in sample separation resulting from sample run order. Similar grouping profiles were observed, with two separate clusters for pre- and post-ST samples. **(b)** Orthogonal partial least squares discriminative analysis (OPLS-DA) of data to maximize separation between the groups.

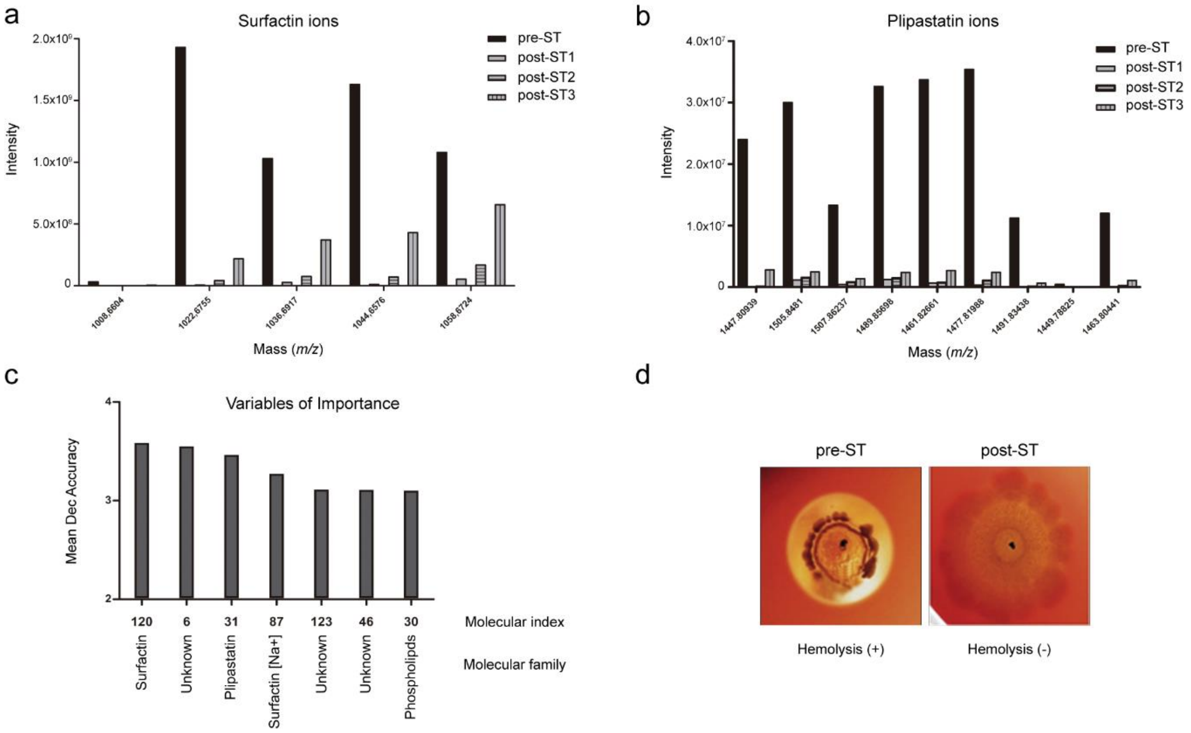


**Supplementary Figure 4. LC-MS/MS-based metabolomic analysis showing chemical signatures that distinguish pre- from post-ST variants**

**(a, b)** Intensities of surfactin and plipastatin ions were strongly reduced in post-ST. **(c)** Random forest analysis indicated that surfactin and plipastatin are major variables determining such separation. **(d)** Loss of hemolytic activity in post-ST variants.

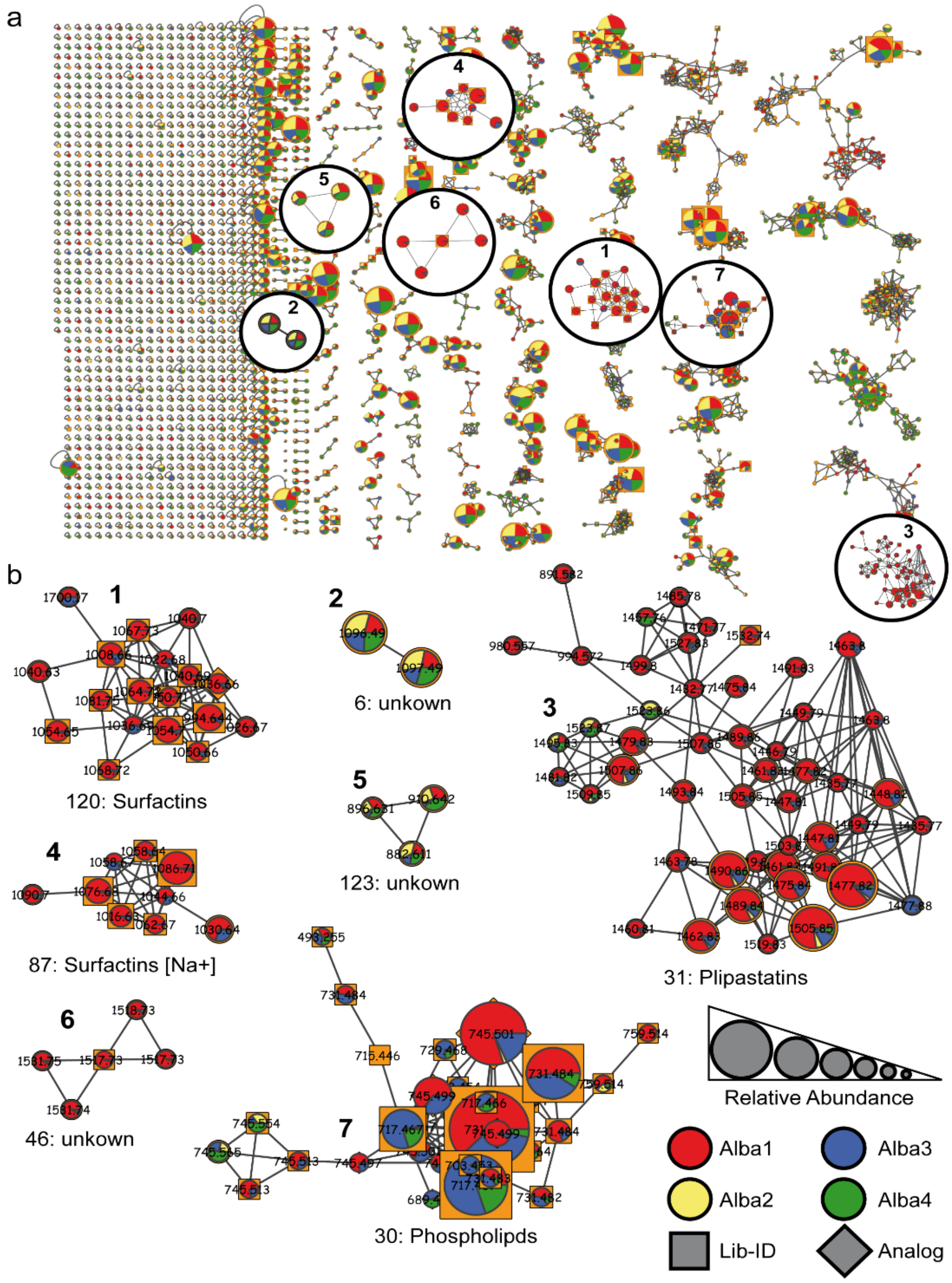


**Supplementary Figure 5. Molecular networks of MS/MS spectra**

**(a**) Global network of all MS/MS spectra (nodes), connected (edges) on the basis of spectral similarity. **(b)** Magnified images of specific subnetworks (molecular families), which correspond to variables of importance that separate groups according to random forest analysis. Pie charts inside the nodes indicate relative feature abundance (averaged XICs, normalized to TIC of particular groups) between groups. Node size indicates relative feature abundance of all samples (averaged XICs, normalized to TIC of all samples).

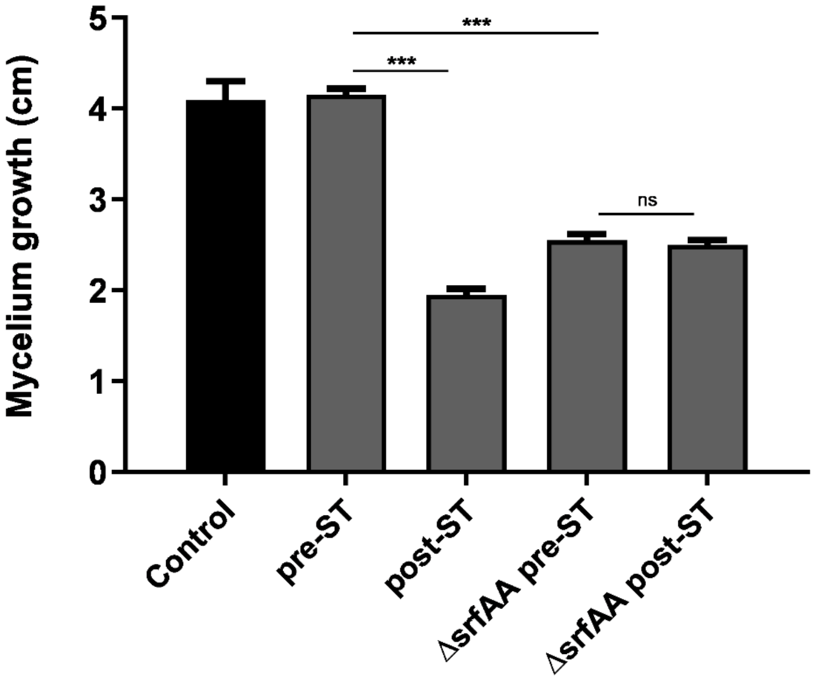


**Supplementary Figure 6. Elimination of surfactin production induces antifungal activity**

Antifungal activities of cell-free supernatants of BsA01 pre- and post-ST and of its Δ*srfAA* mutant were assayed as described in the text. Cell-free supernatants of surfactin-defective mutant Δ*srfAA* and of BsA01 post-ST show clear antifungal activity. No changes in the antifungal activity of Δ*srfAA* were observed before (Δ*srfAA* pre-ST) or after (Δ*srfAA* post-ST) interaction with the fungus. Data shown are mean values of mycelial growth from four independent replicate experiments. ***, significant difference between values ($p < 0.0001$, Tukey's Multiple Comparison Test). ns, no significant difference.

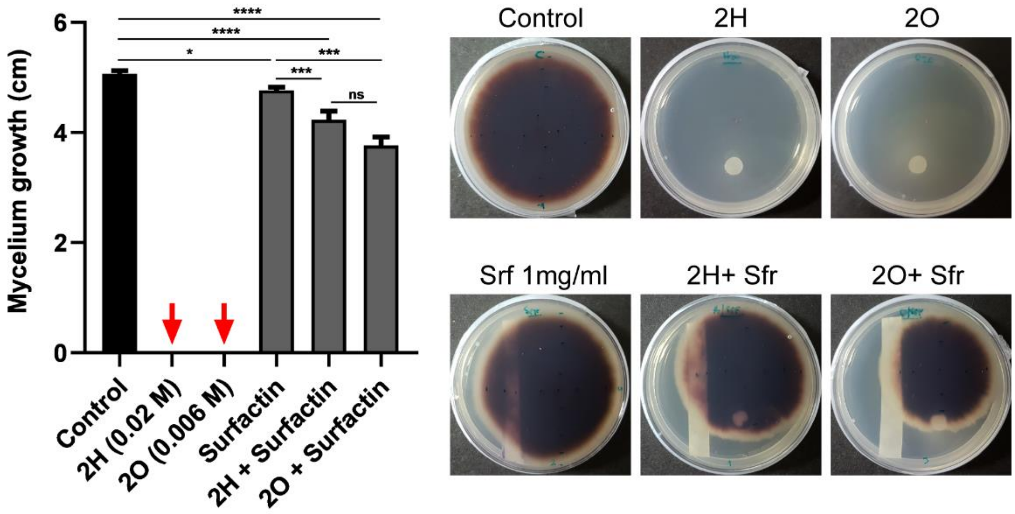


**Supplementary Figure 7. Surfactin interfere with the anti-*S. terrestris* activity of ketones 2-heptanone and 2-octanone**

Suppressor effect of surfactin (1mg/ml imbibed in a filter paper strip with) on the antifungal activity of 2H 0.02 M and 2O 0.006 M, 14 days after inoculation of *S. terrestris*. Data shown are mean values of mycelial growth from three independent replicate experiments; red arrow indicates lethal concentrations of 2-ketones.

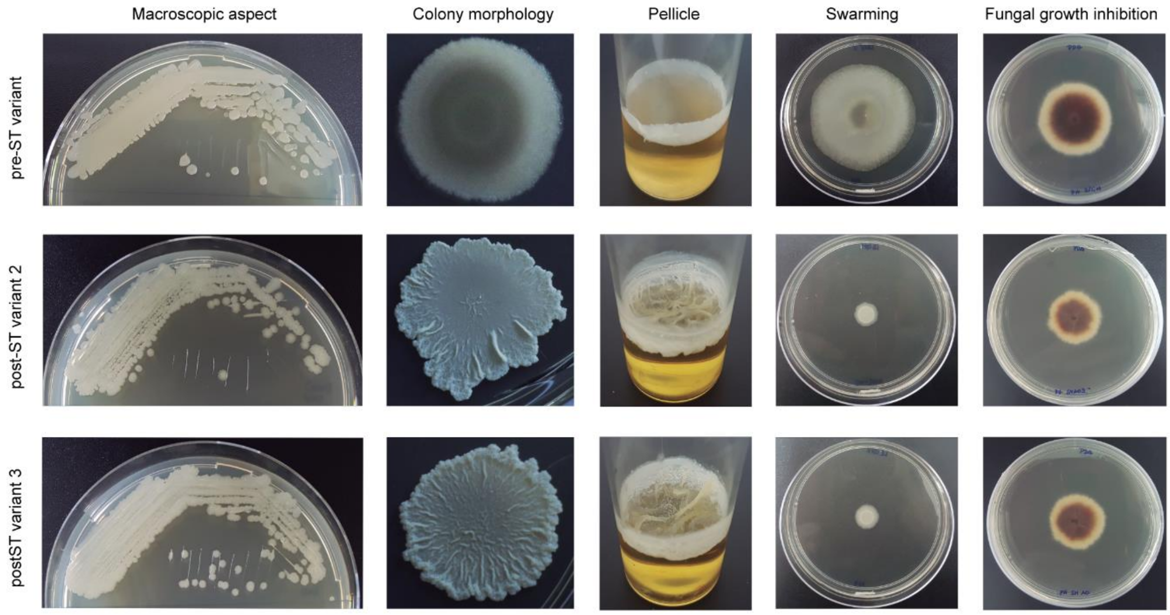


**Supplementary Figure 8. Phenotypic characterization of post-ST variants 2 and 3**

All *B. subtilis* variants obtained following co-culture with *S. terrestris* had similar stable phenotypes. Phenotypes of two other variants whose genomes were sequenced, post-ST variants 3 and 4, are shown here. Similarly to observations of post-ST 1, post-ST 2 and 3 showed rough, wrinkled colony appearance and robust pellicle formation relative to pre-ST. Swarming motility was absent in post-ST 2 and 3. Dishes containing only cell-free supernatants of post-ST 2 and 3 showed strong growth inhibition of *S. terrestris* (7 days after inoculation, at 30°C).

### Supplementary Tables

**Supplementary Table 1.** Comparative whole-genome sequencing analysis of independently obtained post-ST variants, and whole-genome sequencing of pools of individuals (Pool-seq), revealed mutations in *comQXPA* coding regions. The $+1A_{415}$ *comA* insertional mutation was found along with three new mutations: two nonsense substitutions (T215A and C601T) and one 5-nucleotide insertion at position 299. The T215A mutation generated a premature stop codon at position 126 of ComA protein, while the C601T and the 5-nucleotide insertion generated stop codons at 201 and 78, respectively. Two new mutations (both insertions) in *comP* were observed: GC insertion at gene position 1517 generated a truncated version of ComP protein with stop codon at position 512, and AT insertion generated a stop codon at ComP position 232.

| **Variant** | **Scaffold/position** | **Gene** | **Nucleotide substitution** | **Description** |
|---|---|---|---|---|
| Post-ST 1 | 1/69978 | *comA* | Ins A 415 | Transcriptional regulatory protein ComA |
| Post-ST 2 | 1/69978 | *comA* | Ins A 415 | Transcriptional regulatory protein ComA |
| Post-ST 2 | 2/165291 | *yhfV* | ΔA 1089 | Methyl-accepting chemotaxis protein |
| Post-ST 2 | 3/371952 | *spo0A* | G289C | Sporulation two-component response regulator |
| Post-ST 3 | 1/71309 | *comP* | Δ632-731 | Two-component sensor kinase ComP |
| Pool-seq | 1/69979 | *comA* | C601T | Transcriptional regulatory protein ComA |
| Pool-seq | 1/69978 | *comA* | Ins A 415 | |
| Pool-seq | 1/70101 | *comA* | Ins CTGGG 299 | |
| Pool-seq | 1/70185 | *comA* | T215A | |
| Pool-seq | 1/71264 | *comP* | Ins CG 1517 | Two-component sensor kinase ComP |
| Pool-seq | 1/71919 | *comP* | Ins AT 863 | |

**Supplementary Table 2.** Bacterial and fungal strains used in this study.

| Strain | Characteristics | Source |
|---|---|---|
| *Escherichia coli* DH5α | F-Φ80dlacZ ΔM12 minirecA | Our lab |
| *Bacillus subtilis* ALBA01 | Isolate from onion rhizosphere | Our lab |
| *B. subtilis* pre-ST | *B. subtilis* ALBA01 before co-culture with *S. terrestris* | This study |
| *B. subtilis* post-ST | Variants of *B. subtilis* ALBA01 obtained after co-culture with *S. terrestris* | This study |
| *ΔsrfAA* | *B. subtilis* ALBA01 surfactin-defective mutant | This study |
| *ΔsrfAA* pre-ST | Surfactin-defective mutant before co-culture with *S. terrestris* | This study |
| *ΔsrfAA* post-ST | Surfactin-defective mutant after co-culture with *S. terrestris* | This study |
| *Setophoma terrestris* PH06 | Fungal strain isolated from onion rhizosphere | Our lab |

**Supplementary Table 3.** Oligonucleotide primers used in this study.

| Primer | Sequence (5' – 3') | Source |
|---|---|---|
| Fsrfa_nul | CGCGGATCCTGACACGATGTTCAGCCTTC | Modified from [10] |
| Rsrfa_nul | GCGGAATTCCAAAACGGTTTCCTTCGGTA | Modified from [10] |
| FcomA_map | TCAAGCAGCATGATTTCTCG | This study |
| RcomA_map | GTCCGTGAACCGACATTCAG | This study |
| FcomP_map | CGATACGTTTGTATAAAAAGCCAAA | This study |
| RcomP_map | TGTGGATTTTATTTTGAGCAGGT | This study |

**Supplementary Table 4.** MZmine2/ADAP settings used for feature finding.

| | |
|---|---|
| **Mass Detection Module** | |
| Mass Detection | Centroid, Noise level 500 |
| Wavelet transform | Wavelet, Noise level 100 |
| Scale level | 5 |
| Wavelet window size (%) | 0.3 |
| **ADAP Chromatogram Builder Module** | |
| Min group size in # of scans | 5.0 |
| Group intensity threshold | 1000.0 |
| Min highest intensity | 1000.0 |
| m/z tolerance | 0.01 Da/20 ppm |
| **Smooting Module** | |
| Filter width | 25 |
| **Deconvolution Module** | |
| Deconvolution | Savitzky-Golay |
| Min peak height | 5000.0 |
| Peak duration range (min) | 0.01-0.2 |
| Derivative threshold level | 0.1 |
| **ADAP3 Decomposition Module** | |
| Min cluster distance (min) | 0.001 |
| Min cluster size | 8 |
| Min cluster intensity | 50000.0 |
| Find shared peaks | False |
| Min edge-to-height ratio | 0.2 |
| Min delta-to-height ratio | 0.2 |
| Min sharpness | 10.0 |
| Shape-similarity tolerance (0..90) | 80.0 |
| Choice of Model Peak based on | Shaprness |
| **Join Aligner Module** | |
| m/z tolerance | 0.01 Da/20 ppm |
| Weight for m/z | 50.0 |
| Retention time tolerance | 0.05 |
| Weight for RT | 0.05 |
| Require same charge state | False |
| Require same ID | False |
| Isotope m/z tolerance | 0.001-5 |
| **Peak Finder Module** | |
| Intensity tolerance | 0.008 |
| m/z tolerance | 0.01 Da/20 ppm |
| Retention time tolerance | 0.05 |
| RT correction | False |
